# The C99 Fragment Of App Regulates Cholesterol Trafficking

**DOI:** 10.1101/740670

**Authors:** M. Pera, D. Larrea, J. Montesinos, C. Guardia-Laguarta, R.R. Agrawal, K.R. Velasco, Y. Xu, SY Koo, A Snead, A. Sproul, E. Area-Gomez

## Abstract

The link between cholesterol homeostasis and the cleavage of the amyloid precursor protein (APP), and their relationship to the pathogenesis of Alzheimer’s disease (AD) is still unknown. Cellular cholesterol levels are regulated by a crosstalk between the plasma membrane (PM), where most of the cholesterol resides, and the endoplasmic reticulum (ER), where the protein machinery that regulates cholesterol resides. This crosstalk between PM and ER is believed to be regulated by lipid-sensing peptide(s) that can modulate the internalization of extracellular cholesterol and/or its *de novo* synthesis in the ER. Our data here indicates that the 99-aa C-terminal fragment of APP (C99), a cholesterol-binding peptide, regulates cholesterol trafficking between the PM and the ER. In AD models, increases in C99 provoke the upregulation of cholesterol internalization and its delivery to the ER, which in turn result into the loss of lipid homeostasis and the appearance of AD signatures, such as higher production of longer forms of amyloid β. Our data suggest a novel role of C99 as mediator of cholesterol disturbances in AD, and as a potential early hallmark of the disease.

---

The lipid composition of cellular membranes is adjusted constantly to regulate processes such as signal transduction and trans-membrane ionic gradients (*1*). To support these events, a network of enzymes interconnects the metabolism of all lipids, and controls the remodeling of the membrane into functional regions (*2*). Often, these functional regions have the characteristics of lipid-rafts or detergent-resistant domains (*2*). Lipid rafts are transient membrane subregions formed by local increases in cholesterol, which interacts with sphingomyelin (SM) and saturated phospholipids to shield cholesterol from the aqueous phase (*2*). These local elevations in cholesterol create highly ordered membrane microdomains that passively segregate lipid-binding proteins, thereby facilitating protein-protein interaction(s) and the regulation of specific signaling pathways (*2*). The formation of these domains is enabled by “lipid-sensing” proteins with the capacity to bind and cluster cholesterol (*3*). If the lipid-sensing proteins are eliminated from that domain, the lipid rafts are abrogated by hydrolysis of the “shielding” SM via activation of sphingomyelinases (SMases) that convert SM to ceramide (*4*). As opposed to SM, ceramide creates an unfavorable environment for cholesterol (*5*), which can then become accessible for removal via esterification (*4*). Hence, the turnover of lipid-raft domains and their capacity to regulate signaling pathways is tightly linked to the regulation of cholesterol homeostasis. Thus, alterations in the latter would affect the regulation of lipid raft formation, and vice versa.

In the cell, cholesterol is either synthesized *de novo* in the endoplasmic reticulum (ER) or taken up as cholesterol esters (CEs) from lipoproteins (*4*). The *de novo* synthesis of cholesterol is activated by the transport of the sterol regulatory element binding protein isoform 2 (SREBP2) from the ER to the Golgi, and its subsequent activation by proteolytic cleavage (*6*). The processed form of SREBP2 translocates to the nucleus and induces the transcription of cholesterol-synthesizing genes, as well as of the SREBP2 gene itself (*6*). When cholesterol supply is sufficient, SREBP2 is retained in the ER in its uncleaved form, thereby pre-empting the *de novo* synthesis pathway. (*6*).

When taken up from lipoproteins, the internalized CEs are hydrolyzed in endolysosomes to unesterified-free cholesterol, most of which is transferred to the PM (*7, 8*). Once PM cholesterol surpasses a threshold, it is transported to the ER for esterification by the enzyme acyl–coenzyme A:cholesterol acyl-transferase 1 (ACAT1), as a means to detoxify the excess of cholesterol (*7*). The resultant CEs are stored in lipid droplets in the cytosol before being secreted (*7*).

Thus, to preserve cholesterol homeostasis, the cell maintains a crosstalk between the PM, where the bulk of cholesterol resides, and the ER, where the enzymatic activities that ensure cholesterol levels reside (*9*). This crosstalk is believed to be controlled by an as-yet-unknown sensor in the ER that triggers the communication between the PM and the intracellular ER regulatory pool of cholesterol where ACAT1 resides, or MAM (*8, 10*).

The regulation of lipid metabolism is particularly critical within the nervous system (*11*). It is therefore not surprising that lipid deregulation has been described numerous times in neurodegenerative diseases, including Alzheimer’s disease (AD) (*12*). Specifically, cholesterol anomalies in AD have been widely reported (*12*), but the field currently lacks consensus as to their cause(s).

The “amyloid cascade hypothesis” in AD pathogenesis states that increases in the levels of β-amyloid peptide (Aβ), derived from APP processing, trigger neurodegeneration (*13*). These higher levels of Aβ in AD are the consequence of corresponding increases in the cleavage of endocytosed full-length APP by β-secretase to produce the immediate precursor of Aβ, the 99-aa C-terminal domain of APP (C99), and its subsequent processing by γ-secretase (*13*). These alterations in APP metabolism are due to mutations in the *PSEN1* [presenilin-1 (PS1)], *PSEN2* [presenilin-2 (PS2)], and *APP* genes in familial AD (FAD), or by unknown causes in sporadic cases (SAD) (*13*). Further linking AD and cholesterol, APP-CTFs processing occurs in lipid rafts (*14*). Notably, elevations in C99 have been shown to contribute to AD (*15*) causing endosomal dysfunction (*16*), and hippocampal degeneration (*17*).

Previously, we and others found that C99, when delivered to the ER for cleavage by γ-secretase, is not distributed in the ER homogeneously, but concentrated at mitochondria-associated ER membranes (MAMs) (*18–20*). MAM is a lipid raft in the ER (*21, 22*), involved in the regulation of lipid homeostasis (*23*). We showed that in AD cell and animal models, there is an increased of C99 at MAM (*20*), resulting in the upregulation of MAM activities (*22, 24*), including SMases, and cholesterol esterification by ACAT1 (*20*). Remarkably, inhibition of C99 production caused the inactivation of these MAM functions (*20*).

We now report that, by means of its affinity for cholesterol (*25*), the pathogenic buildup of C99 in the ER (*20*) induces the uptake of extracellular cholesterol and its trafficking from the PM towards the ER, resulting in the persistent formation, activation and turnover of MAM domains. Altogether, our data suggest a pathogenic role for elevations in C99 via upregulation of cholesterol trafficking and MAM activity, which in turn abrogate cellular lipid homeostasis and disrupt the lipid composition of cellular membranes.

## RESULTS

### Deficiency in γ-secretase activity triggers cellular cholesterol uptake and trafficking to MAM

Co-activation of SMase(s) and ACAT1 is a mechanism by which cells “detoxify” membranes from an excess of cholesterol (*10*). Therefore, we hypothesized that the upregulation of sphingolipid turnover and of cholesterol esterification caused by increases in uncleaved C99 (*20*), could be the consequence of the activation of this detoxifying mechanism, provoked by elevations in cholesterol in cell membranes.

To test this, we measured the concentration of cholesterol in membranes and subcellular fractions from mouse embryonic fibroblasts (MEFs) null for both *PSEN1* and *PSEN2* (PS-DKO) (*26*) (Fig. 1A) and from homogenates from AD fibroblasts (Fig. S1A) by liquid chromatography-mass spectrometry (LC-MS) (*27*), and found that they displayed increased levels of free cholesterol when compared to controls (Fig. 1A). This increase in membrane-bound cholesterol was highly significant in ER and MAM membranes, which, in light of previous data (*22*), suggests that the upregulation of cholesterol esterification in AD cell models is the result of cholesterol buildup in membranes and subsequent elimination by esterification. We were able to recapitulate this increase in free cholesterol levels in MEFs in which both *APP* and its paralog *APLP2* were knocked out (APP-DKO) (*28*), transiently transfected with a plasmid expressing C99, compared to controls (Fig. 1B). Conversely, APP-DKO cells expressing either the APP-C83 peptide (produced by the cleavage of APP by α-secretase) or incubated with amyloid oligomers did not show these cholesterol elevations, suggesting that C99, and no other APP fragment, affected cholesterol homeostasis (Fig. 1B).

**Figure 1.**
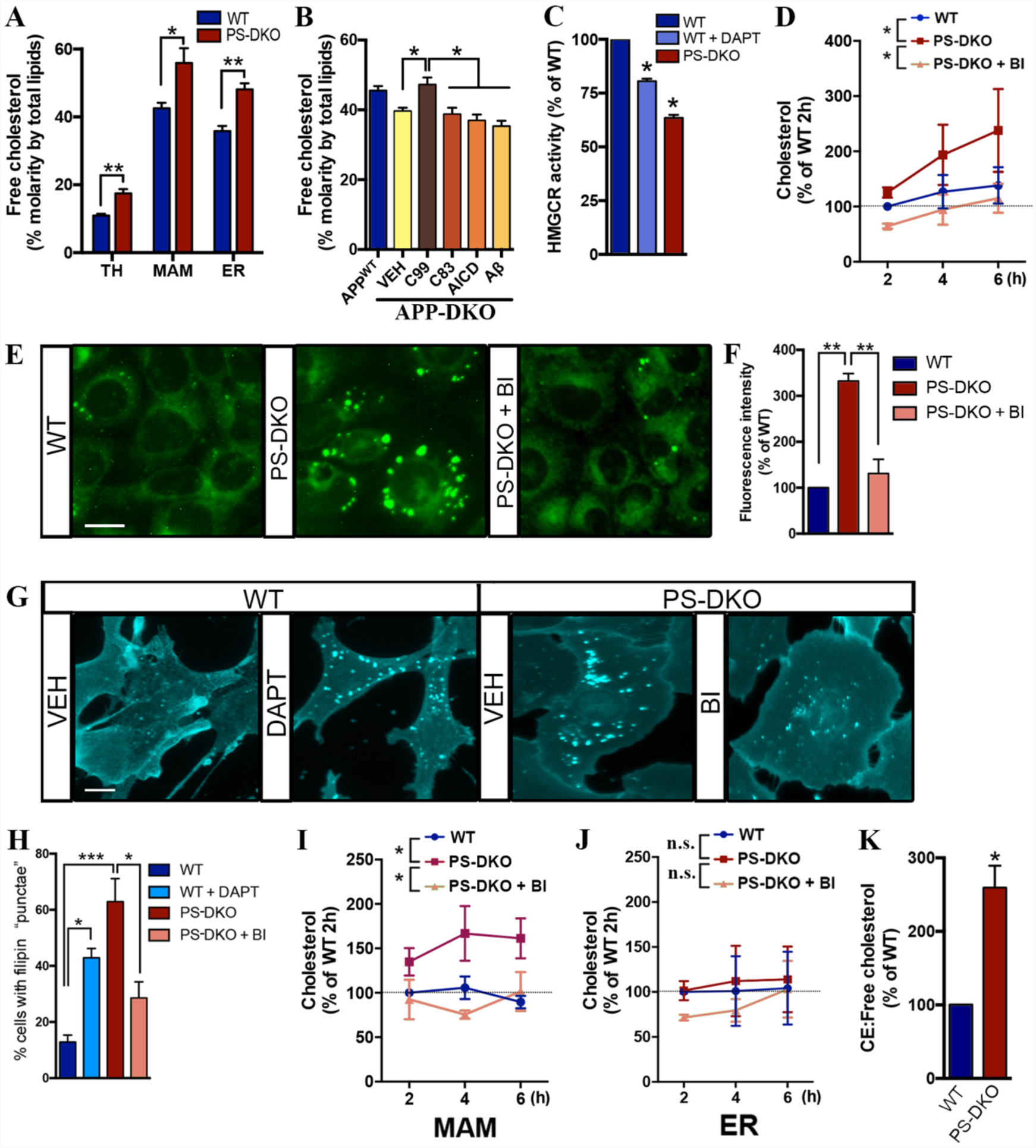
C99 accumulation promotes cholesterol uptake and trafficking to MAM. **(A)** Cholesterol levels are increased in PS-DKO cells when compared to controls. Levels of cholesterol were quantified by lipidomics analysis in total homogenates (TH), ER or MAM fractions from each condition. Lipid units are represented as molar mass over total moles of lipids analyzed (mol%) (* p< 0.05, ** p <0.01, unpaired t-test vs WT, n = 3). **(B)** Cholesterol levels assessed by lipidomics in total homogenates of APP-DKO cells transiently expressing C99 were higher when compared to the same cells expressing C83 or AICD constructs or treated with 5 μM Aβ_42_ oligomers for 16h. Cholesterol levels in APP-WT cells are shown as a control. Lipid units are represented as molar mass over total moles of lipids analyzed (mol%). One-way ANOVA: F_(5,18)_ = 8.63, p < 0.001. (* p<0.05 vs APP-DKO, n = 4). **(C)** Measurement of HMGCR enzymatic activity showed a decrease rate of *de novo* synthesis of cholesterol in PS-DKO cells and in WT cells treated with DAPT (* p< 0.05, one-sample t-test vs WT, n = 3). **(D)** Increased uptake of ^3^H-cholesterol in PS-DKO cells compared to WT or PS-DKO cells treated with BACE inhibitor, BI. (n = 3 independent experiments). Two-way repeated measures ANOVA (Time, Group): Group: F_(2,4)_ = 7.94, p = 0.04, η=0.29. (* p< 0.05 vs PS-DKO). **(E)** Uptake of fluorescently-labeled cholesterol in WT and PS-DKO cells +/-BACE inhibitor (BI). Scale bar = 20 μm. **(F)** Fluorescence intensity was calculated by ImageJ analysis (30-50 cells/condition from at least 3 independent experiments). One-way ANOVA: F_(2,6)_ = 38.75, p < 0.001. (** p<0.01 vs PS-DKO). **(G)** Cholesterol internalization was detected by filipin staining upon the indicated treatments in WT or PS-DKO MEFs. Scale bar = 20 μm. Quantification by ImageJ analysis is shown in **(H)** (30-50 cells/condition from at least 3 independent experiments). One-way ANOVA: F_(3,8)_ = 15.28, p < 0.01. (* p<0.05, *** p<0.001). **(I-J)** Control and PS-DKO cells treated with BI were incubated with ^3^H-cholesterol for the indicated times. Subsequent subcellular fractionation showed that C99 accumulation in PS-DKO promotes cholesterol uptake and trafficking to MAM compared to WT and BI-treated PS-DKO cells. **(I)** MAM and **(J)** ER in WT and PS-DKO cells +/-BI (n = 3 independent experiments). Two-way repeated measures ANOVA (Time, Group) for MAM; Group: F_(2,4)_ = 11.39, p = 0.022, η=0.52; Group: F_(3,36)_ =10.45, p < 0.0001, η=0.27 (* p< 0.05 vs PS-DKO). Two-way ANOVA for ER was not significant. **(K)** Ratio of CE:unesterified cholesterol measured by lipidomics analysis of homogenates from WT and PS-DKO cells. (* p<0.05, one-sample t-test vs WT, n = 3).

To determine whether cholesterol increases in AD cells occurred via upregulation of the *de novo* cholesterol synthesis, we quantified the activity of the 3-hydroxy-3-methylglutaryl-CoA reductase (HMGCR), the rate-limiting enzyme in the synthesis of cholesterol, in γ-secretase-deficient cells and in controls. HMGCR activity was reduced significantly in PS-DKO MEFs, in cells treated with γ-secretase inhibitors (DAPT) (Fig. 1C), and in AD fibroblasts (Fig. S1B), in agreement with previous observations (*29*). Consistently, total levels of SREBP2 were significantly lower in the PS-DKO cells, as was that of mature SREPB2 after DAPT treatment (Fig. S1C).

Given this reduced *de novo* synthesis of cholesterol, we then measured the rate of uptake of extracellular cholesterol in PS-DKO cells and controls, by incubating them with ^3^H-cholesterol and following its internalization over time. Mutant cells showed an enhanced rate of cholesterol uptake compared to controls, whereas treatment of these cells with a BACE1 inhibitor had the opposite effect (Fig. 1D). We also observed increased cholesterol uptake in neuronal cells silenced for PS1 alone or for both PS1+PS2 (Fig. S1D), and in AD fibroblasts (Fig. S1E).

Supporting this result, the uptake of fluorescently-labeled cholesterol was also increased in PS-DKO cells compared to controls (Fig. 1E-F). Interestingly, this increase in either radioactive (Fig. 1D) or fluorescently-labeled (Fig. 1E-F) cholesterol uptake was abrogated upon BACE1 inhibition, which indeed, suggests a role for C99 in the regulation of cholesterol internalization.

Staining of our cells with filipin, a dye that binds to free cholesterol, showed that the *distribution* of free cholesterol was also markedly different in PS-DKO mutant (Fig. 1G-H), and in AD fibroblasts vs. controls (Fig. S1F), which showed a higher degree of cholesterol *punctae* in the cytosol, suggesting an increased rate of cholesterol influx into mutant cells.

To address the subcellular destination of the internalized cholesterol, we tracked the uptake of ^3^H-cholesterol, and its delivery to MAM and bulk ER by pulse-chase analysis and subcellular fractionation. We found that in PS-DKO cells the *rate* of cholesterol incorporation into MAM was higher when compared to controls, and was abrogated upon BACE1 inhibition (Fig. 1I-J). We were able to recapitulate this phenotype (Fig. S1G-H) in iPSCs in which a pathogenic mutation in APP (London mutation; APP^V717I^) was knocked into both alleles using CRISPR/Cas9, which showed higher levels of C99 relative to isogenic control iPSCs (Fig. S1J). This enhanced trafficking of cholesterol from the PM to the ER-MAM was also reflected in an increased ratio of cholesteryl esters: free cholesterol (CE:FC) (*8, 30*) in PS-DKO cells (Fig. 1K), in our cell models expressing mutant APP^V717I^ (Fig. S1I), and in cells from AD patients (Fig. S1K).

In cellular membranes, a pool of cholesterol in the PM is complexed with SM, preventing its mobilization (*7, 31*), although treatment with SMases can release the membrane-bound cholesterol and induce its transport to the ER (*30*). Our previous data revealed that increases in MAM-localized C99 trigger the upregulation of SMase(s) activity (*20*). Thus, we asked whether the increases in cholesterol trafficking towards ER-MAM domains were a consequence of sustained SMase activity provoked by increases in MAM-C99.

To test this, we treated our cells with SMase inhibitors, and then stained them with filipin or LipidTox Green, a dye that visualizes lipid droplets (LDs). As before (*20*), incubation with the SMase(s) inhibitors reduced cholesterol esterification by ACAT1 and the accumulation of LDs in PS-DKO cells (Fig. 2A-B), in DAPT-treated SH-SY5Y cells and AD patient fibroblasts (Fig. S2). Thus, the increase in ACAT1 activity and in LD production associated with elevated C99 was facilitated by activation of SMase(s), consistent with previous findings (*30*). However, cholesterol uptake was not reversed by SMase inhibition in either DAPT-treated WT cells (Fig. 2C-D), or in PS-DKO cells (Fig. 2E-F), thereby indicating that SMase upregulation is likely not a cause, but rather a consequence, of a prior enrichment of cholesterol in membranes.

**Figure 2.**
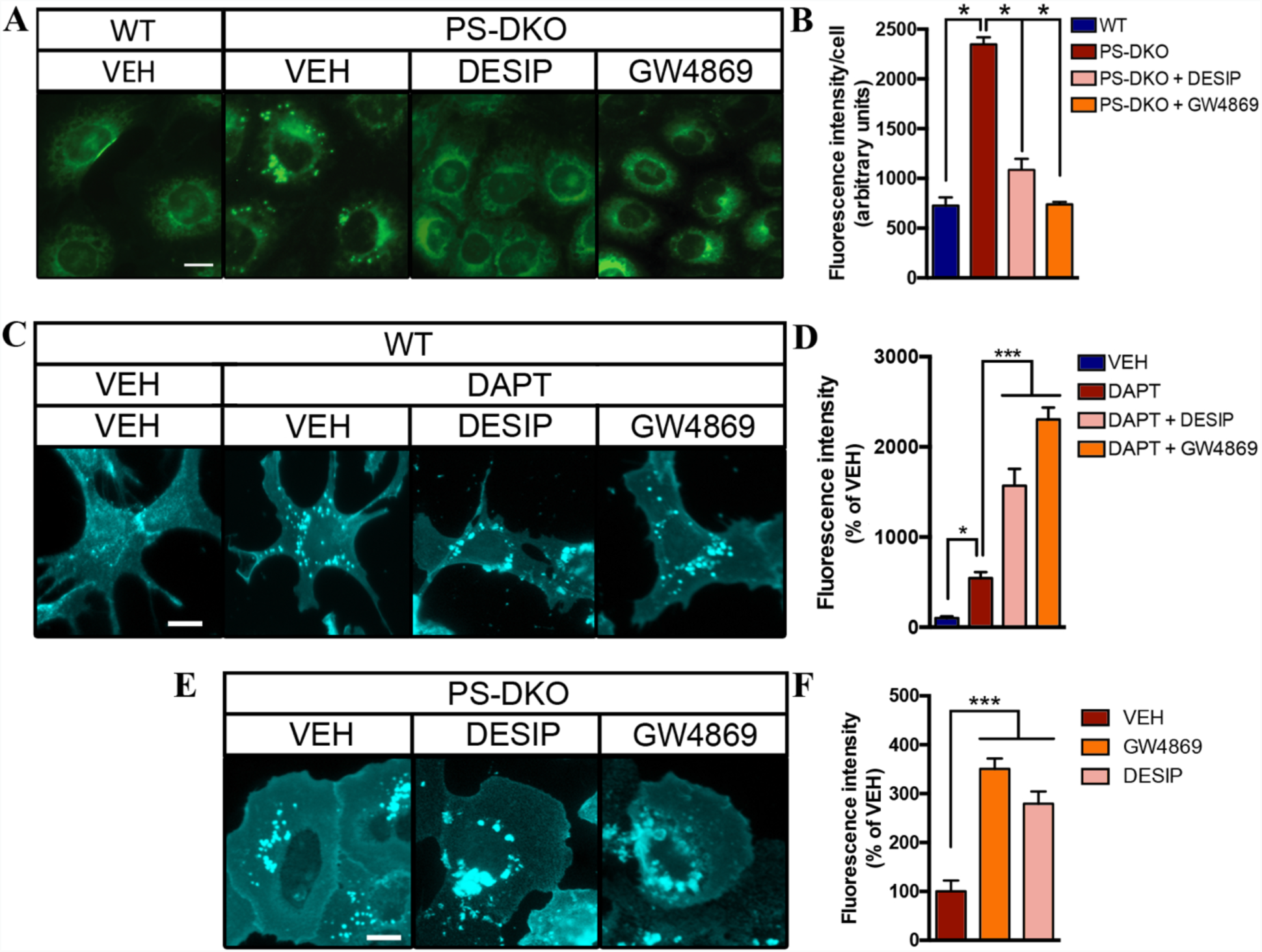
SMase inhibition prevents LDs generation but fails to correct the increase in cholesterol uptake caused by C99 accumulation. Lipid droplet visualization in **(A-B)** WT and PS-DKO MEFs +/-SMase inhibitors (10 μM desipramine or 5 μM GW4869) for 12-16h or DMSO (VEH) and stained with Lipidtox Green to detect lipid droplets. Quantification by ImageJ analysis is shown in **B** (30-50 cells/condition from at least 3 independent experiments). One-way ANOVA: F_(3,8)_ = 92.02, p<0.001. * p< 0.05. Scale bar = 20 μm. **(C-F)** Filipin staining of WT **(C)** or PS-DKO **(E)** under the indicated treatments. Note how SMase inhibitors do not inhibit cholesterol internalization, as measured by increases in the amount of filipin-*punctae*, in DAPT-treated WT or PS-DKO cells, supporting the idea that SMase activation is a detoxifying mechanism activated to balance the increased uptake of extracellular cholesterol. Quantification by ImageJ for WT cells is shown in **D** (One-way ANOVA: F_(3,19)_ =71.91, p < 0.001, n=5-7); and for PS-DKO cells in **F** (One-way ANOVA: F_(2,15)_ =31.81, p < 0.001, n=6). * p<0.05, *** p<0.001. Scale bar = 20 μm.

Taken together, our results suggest that γ-secretase-deficient cells suffer from an excessive internalization of cholesterol and delivery to MAM, where its esterification by ACAT1 is facilitated by SM hydrolysis by SMase(s)(*20*). Our data also indicate that this mobilization of cholesterol to ER-MAM has the further effect of reducing *de novo* cholesterol biosynthesis via inactivation of SREBP2.

### Localization of C99 at MAM depends on its cholesterol binding domain

The C99 APP fragment contains a cholesterol binding motif within its transmembrane domain (*25*), that could promote its localization to lipid rafts (*32*), such as MAM (*20, 22*). To test whether C99’s cholesterol binding domain (CBD) was necessary for its localization to MAM, we transfected APP-DKO cells with plasmids expressing WT C99 and a mutant C99 construct with reduced affinity for cholesterol (G_700_AII_703_G_704_ was mutated to A_700_AIA_703_A_704_; named C99^CBD^ for simplicity), and treated them with DAPT to impede C99 cleavage before analyzing its subcellular localization (Fig. S3A) by confocal analysis of these APP-DKO cells also expressing mitochondria (MitoDsRed) and ER (Sec 61β-BFP) markers (Figs. 3A, 5A, S3 and S5A). While C99^WT^ showed a perinuclear pattern of colocalization with ER and mitochondria, C99^CBD^ presented a less marked perinuclear localization and a decreased association with mitochondria (Figs. 5A, 5B, and S5A).

**Figure 3.**
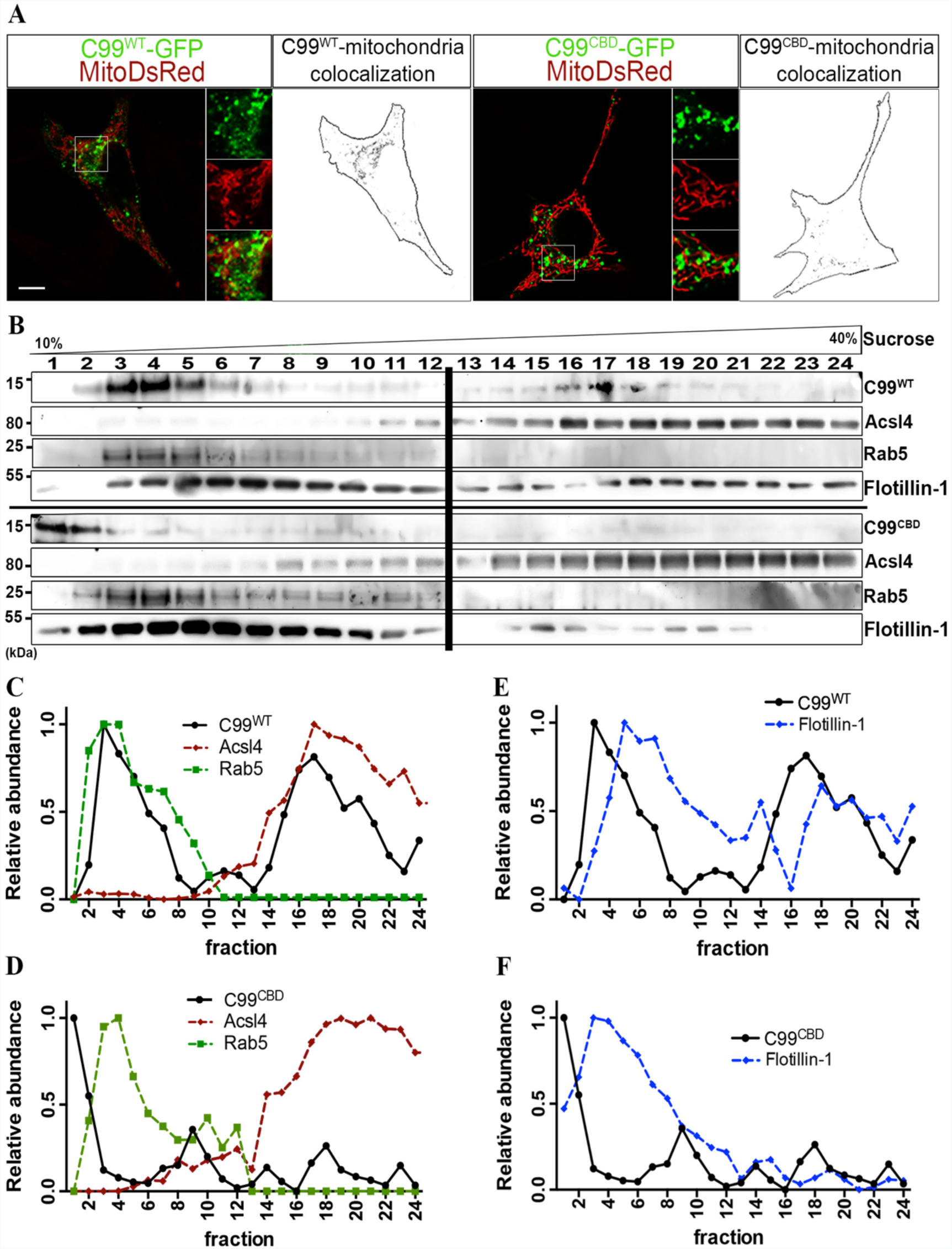
The cholesterol binding domain of C99 is necessary for its localization to MAM. **(A)** Confocal images showing mitochondria (MitoDsRed, in red) and the distinct distribution of GFP-tagged C99^WT^ or C99^CBD^ (in green). Scale bar = 10 μm. Insets show 5× amplifications of individual (green and red) and merged images. Black and white panels show C99-mitochondria colocalization. **(B)** Crude membrane fractions from APP^DKO^ cells expressing C99^WT^ or C99^CBD^ were treated with Triton-X and purified on a continuous sucrose gradient for 16h. Fractions from the gradients were analyzed by western blot for the indicated proteins [two parallel gels (bold line)]. One representative experiment of 5 total trials is shown. Relative abundance along the gradient for each protein is shown as indicated **(C-F)**.

To gain insight into these changes in C99 localization, we run membrane homogenates from these same transfected cells through a continuous density sucrose gradient, and analyzed the migration of specific markers by western blot (Fig. 3B). As shown before (*20*), C99^WT^ comigrated with endosomal (Rab5) and MAM markers (Acsl4) (Fig 3C). However, C99^CBD^ showed reduced comigration with MAM markers (Fig. 3D). As expected, flotillin, a protein with affinity for cholesterol, also comigrated with MAM markers in cells transfected with C99^WT^ (Fig. 3E). Conversely, in the presence of C99^CBD^, flotillin showed a reduced comigration with MAM proteins (Fig. 3F), suggesting that C99 binding to cholesterol not only is important for its localization to MAM, but for the proper formation of MAM itself.

As a lipid raft, MAM (*22*), is a transient functional domain formed by local increases in cholesterol mediated by peptides with CBD capable of docking cholesterol until it coalesces into a rigid domain (*3*). Thus, in light of our results, we hypothesized that C99, when delivered to the ER, docks to cholesterol via its CBD, thereby helping form MAM domains. To test this, we decided to analyze C99 affinity for cholesterol using a PhotoClick-cholesterol chemistry approach (Fig. 4A), a photoreactive alkyne cholesterol analog that faithfully mimics native cholesterol, and that can serve as a tool to determine protein affinity for cholesterol (*33*). We incubated APP-DKO cells transiently expressing C99^WT^ or C99^CBD^ with this *trans*-sterol probe. Subcellular fractions from these cell models were then conjugated to an azide-biotin tag by click chemistry, followed by a pull-down assay using streptavidin beads (Figs. 4A and S4A-B). As a proof of principle, we were able to detect increased levels of C99^WT^ bound to cholesterol in MAM fractions from PS-DKO cells (Fig. S4C).

**Figure 4.**
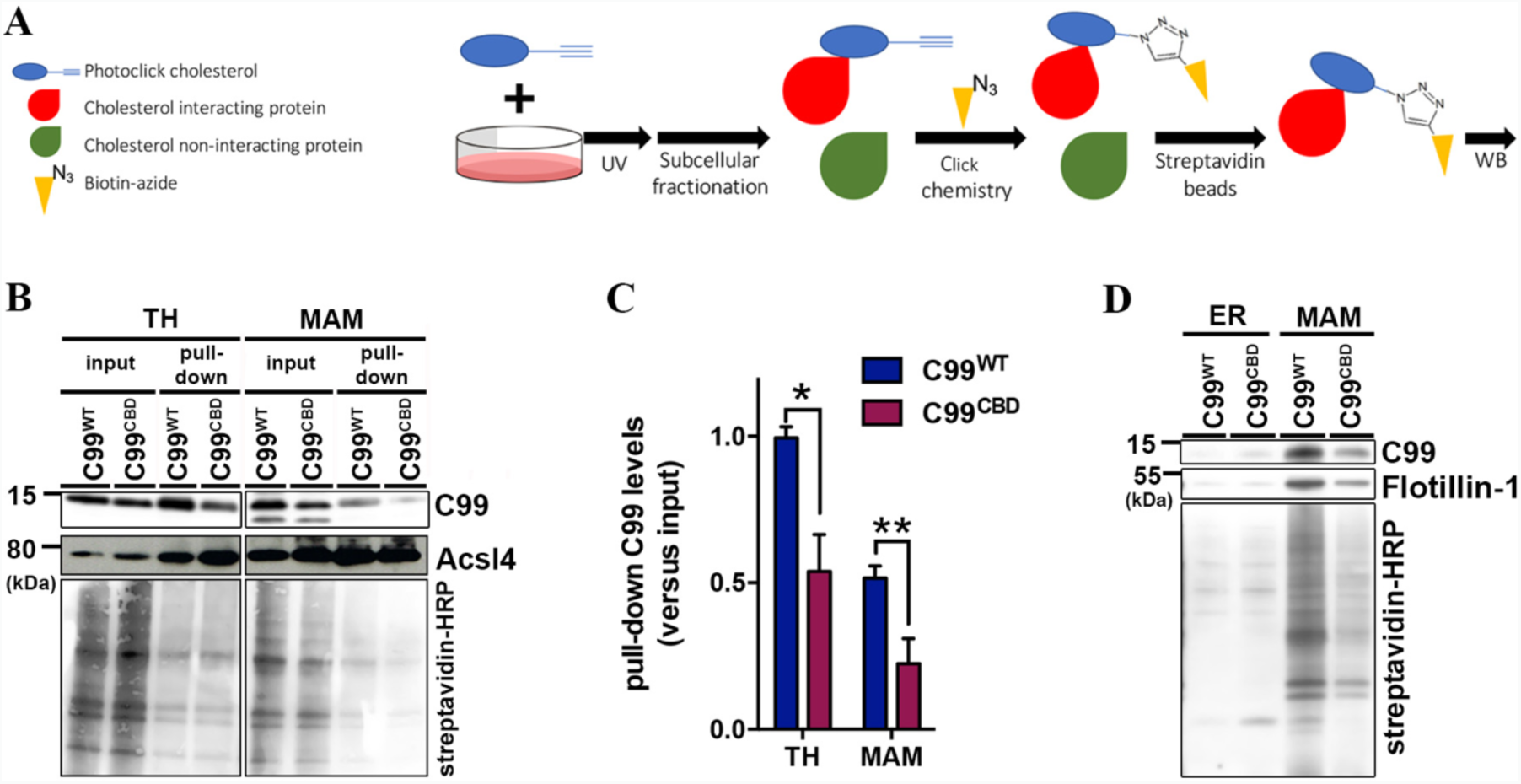
C99’s cholesterol binding domain is necessary for cholesterol trafficking to MAM domains. **(A)** Diagram of PhotoClick cholesterol methodology to detect C99 interaction with cholesterol. **(B)** Representative immunoblot of C99 levels in TH and MAM fractions of APP-DKO cells expressing C99^WT^ or C99^CBD^ before and after cholesterol pull-down. Acsl4 was used as a MAM marker. Streptavidin-HRP was used to detect total biotinylation (biotin conjugated to PhotoClick cholesterol). **(C)** Quantification (n=5) of pulled-down C99 levels versus input. Two-way repeated measures ANOVA (Fraction, CBD): Fraction: F_(1,4)_ =14.43, p < 0.05, η=0.38; CBD: F_(1,4)_=36.41, p < 0.01, η=0.34. * p<0.05, ** p<0.01. **(D)** Immunoblot showing the levels of pulled-down PhotoClick cholesterol (streptavidin-HRP) in ER vs MAM. Note how the levels of pulled-down C99 in the ER are negligible when compared to those from MAM.

The levels of click-cholesterol in homogenates of APP-DKO cells expressing C99^WT^ or C99^CBD^ were comparable (Fig. 4B-C). However, pull-down of the added PhotoClick-cholesterol revealed that the amount of C99^CBD^ bound to this lipid was significantly reduced when compared to that of C99^WT^ (Fig. 4B-C), demonstrating that, as reported (*25*), the C99 residues G_700_AII_703_G_704_ are pivotal for C99 cholesterol binding.

Moreover, cells transfected with C99^CBD^ exhibited significant reductions in the amount of click-cholesterol that was transferred to MAM (Fig. 4B). Consistently, lipidomics analysis showed reductions in cholesterol in MAM fractions from C99^CBD^ vs C99^WT^ cells (Fig. S4D). Similar to the results obtained in homogenates, pull-down of the click cholesterol present in MAM fractions from cells transfected with C99^CBD^ displayed lower levels of bound C99 (Fig. 4B-C-D).

Taken together, our results support the idea that the cholesterol binding domain of C99 is necessary to induce cholesterol internalization for the formation of MAM in the ER (*34*). In support of this conclusion, the apposition between ER and mitochondria was significantly decreased in APP-DKO cells expressing C99^CBD^ when compared to controls (Fig. 5A-B).

**Figure 5.**
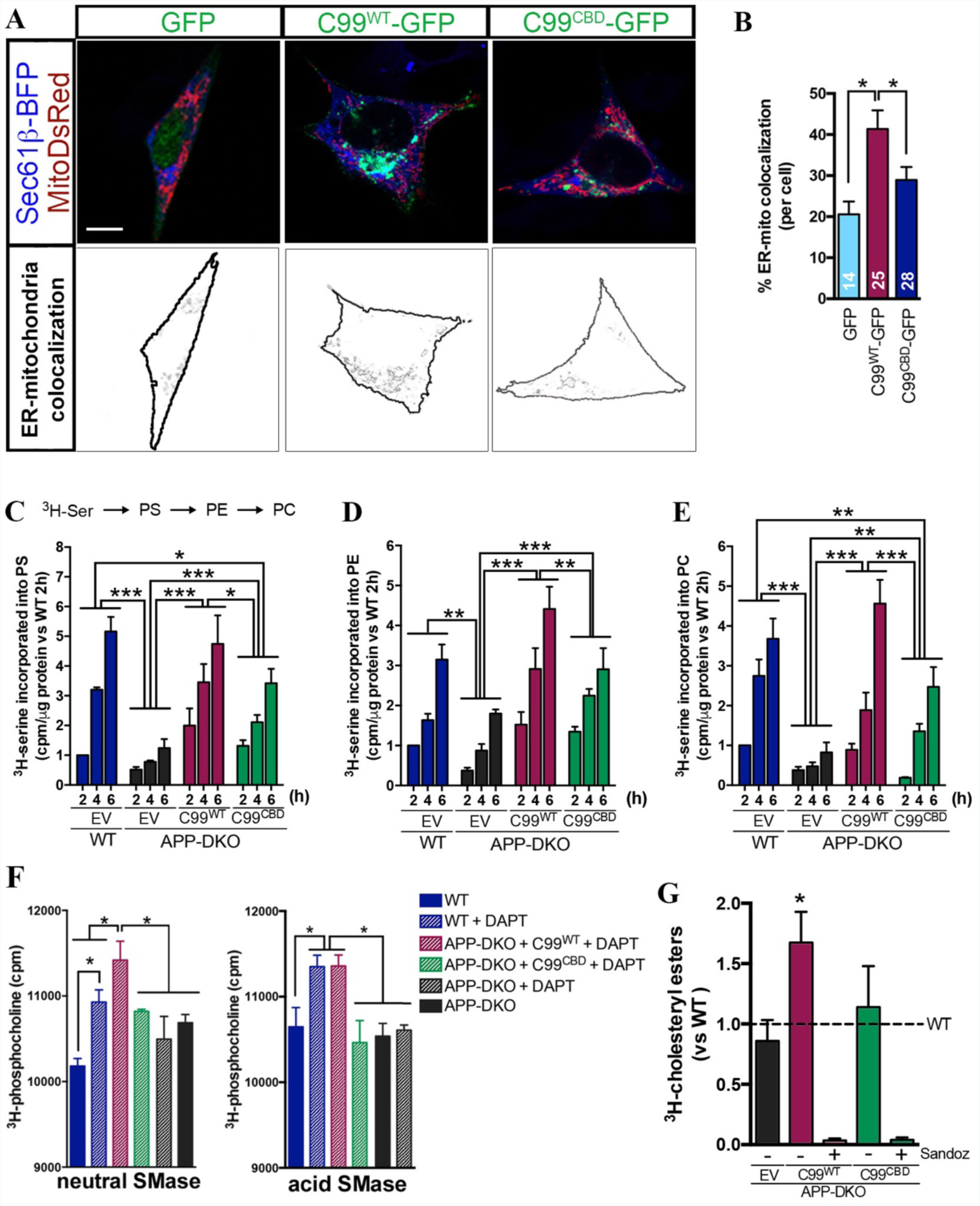
C99 CBD facilitates MAM formation and activation. **(A)** ER-mitochondria apposition was assessed by measuring co-localization of ER signal (Sec61β-BFP, in blue) and mitochondria signal (MitoDsRed, in red) in positively transfected cells (in green, GFP, GFP-tagged C99^WT^ or GFP-tagged C99^CBD^). Representative confocal images are shown in the upper panel and ER-mitochondria colocalization in the lower panel. Scale bar = 10 μm. Single red, blue and green channels are shown in Fig. S5A. **(B)** Percentage of ER-mitochondria co-localization per cell. The number of cells analyzed, from 3 different experiments, is shown as a white number in each column. One-way ANOVA: F_(2,64)_ = 6,284, p < 0.01. * p<0.05. **(C-G)** The effect of C99^CBD^ compared to C99^WT^ on MAM functionality was analyzed using different assays: **(C-E)** Pulse-chase analysis of phospholipid transfer in the different groups was analyzed by labelling with ^3^H-serine for the indicated times (h) and the levels of **(C)** ^3^H-phosphatidylserine, PS, **(D)** ^3^H-phosphatidylethanolamine, PE, and **(E)** ^3^H-phosphatidylcholine, PC, were assessed by TLC (n=3). EV, empty vector. Two-way ANOVA (Time, Group). <u>For PS</u>: Time: F_(2,24)_ = 30.93, p < 0.001, η=0.38; Group: F_(3,24)_ =20.58, p < 0.001, η=0.38; Time x Group: F_(6,24)_ = 2.69, p < 0.05, η=0.10. <u>For PE</u>: Time: F_(2,24)_ = 40.19, p< 0.001, η=0.48; Group: F_(3,24)_ =18.83, p < 0.001, η=0.33. <u>For PC</u>: Time: F_(2,24)_ = 45.51, p < 0.001, η=0.43; Group: F_(3,24_) =22.28, p < 0.001, η=0.32; Time x Group: F_(6,24)_ = 4.81, p < 0.05, η=0.14. (* p<0.05, ** p<0.01, *** p<0.001). **(F)** SMase activity. Activity of neutral and acid SMase in the indicated conditions was assessed measuring conversion of ^3^H-sphingomyelin into 3H-phosphocholine. One-way ANOVA; For neutral SMase: F_(5,12)_ = 6.36, p < 0.01. For acid SMase: F_(5,12)_ = 5.79, p < 0.01 (representative experiment with 3 technical replicates of 3 independent experiments). * p<0.05. **(G)** Cholesterol esterification by ACAT1. Levels of ^3^H-cholesterol incorporated into ^3^H-cholesteryl esters were analyzed by TLC. Dashed line indicates APP-WT levels used as control. Treatment with Sandoz 58-035, a specific ACAT1 inhibitor, caused a ~95% reduction in cholesterol esterification (n=3; * p<0.05, one-sample t-test vs APP-WT). EV, empty vector. In Figures C, D, E and G, all the groups were incubated with DAPT.

We measured MAM activity in APP-DKO cells expressing either C99^WT^ or C99^CBD^ vs. controls by analyzing the synthesis and transfer of phospholipids between ER and mitochondria (*23*). As shown before (*20*), overexpression of C99^WT^ resulted in significant increases in the crosstalk between ER and mitochondria (Fig. 5C-E). Conversely, expression of C99^CBD^ showed significantly lower effects on MAM activity, thus corroborating the importance of C99 cholesterol binding on MAM activation. Consistently, C99^WT^ expression in the presence of DAPT resulted in the upregulation of SMase activity (Fig. 5F), and of cholesterol esterification by ACAT1 (Fig. 5G), whereas expression of C99^CBD^ did not show any difference when compared to controls. In agreement with these data, APP-DKO expressing C99^WT^ showed significant increases in LDs (Fig. S5A-B), and a higher ratio of CE:FC (*30*) (*10*) when compared to its C99^CBD^ counterpart and controls (Fig. S5C).

Altogether, our results suggest that C99 capacity to recruit and cluster cholesterol in the ER triggers the formation and activation of MAM domains. Moreover, the increased concentration of uncleaved C99 fragments in the ER in cells from AD patients induces the constant turnover of MAM domains by activating the entry of cholesterol and its trafficking to MAM (Fig. 6).

**Figure 6.**
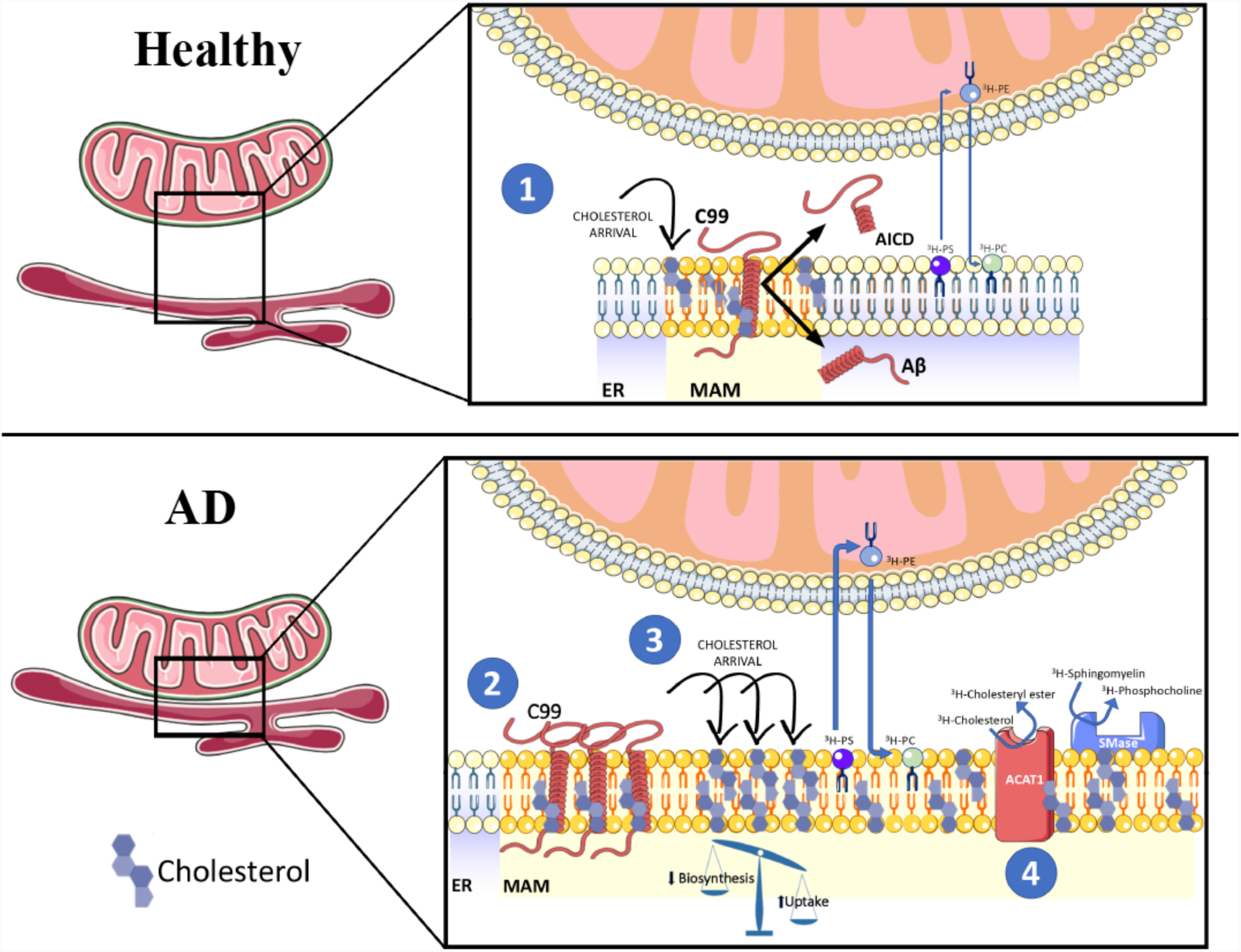
Schematic representation of the potential role of C99 in the regulation of cholesterol trafficking, and its relevance to AD. By means of its affinity for cholesterol, uncleaved C99 at the ER, induces the uptake and retrograde transport of cholesterol from the PM to the ER, resulting in the formation of a lipid raft domain, or MAM. These C99-dependent lipid-rafts would passively segregate and organize lipid-binding proteins, thereby facilitating their interaction and the regulation of specific signaling pathways. Failure to cleave C99 completely would result in a futile cycle of continuous uptake of extracellular cholesterol and its mobilization from the PM to ER, resulting in the upregulation of MAM formation and activation, which in turn would cause the upregulation of SMases, ACAT activity, and LD deposition. Closing the cycle, this accumulation of cholesterol in membranes also induces APP internalization and its interaction and cleavage by BACE1, and the downregulation of α-secretase activity

## DISCUSSION

In previous reports, we and others have found that, γ-secretase activity is localized in the ER, although not homogeneously; rather it is enriched in MAM domains (*18, 19*). Moreover, alterations in γ-secretase activity result in the functional upregulation of this ER region and in increased ER-mitochondria apposition (*22, 24*).

Recently, we showed that the γ-secretase substrate, C99, is also enriched in MAM and that, alterations in γ-secretase activity provoke an accumulation of this APP fragment in MAM membranes that in turn, cause the upregulation of MAM activities, increased cholesterol esterification and sphingolipid turnover, and mitochondrial dysfunction (*20*). We now show that these phenotypes are consequences of the continuous uptake of extracellular cholesterol and its delivery to the ER, provoked by increased levels of C99.

Our data supports a model (Fig. 6) in which C99 at MAM, induces the uptake and transport of cholesterol from the PM to the ER (*9*), via an as-yet unknown mechanism, which results in the inhibition of the SREBP2-regulated pathway(s) (*6*). We propose that, under normal circumstances (Fig. 6, **upper panel**), C99, via its CBD, causes cholesterol to cluster in the ER, resulting in the formation of MAM domains. As this MAM-cholesterol pool expands in, it will activate the hydrolysis of sphingomyelin by SMases, exposing cholesterol to ACAT1 for esterification, which will decrease the concentration of membrane-bound cholesterol, resulting in the dissolution of the lipid raft (*4*). In this way, C99 promotes a self-regulating feedback loop to help maintain intracellular cholesterol levels.

Under this point of view, the failure to cleave C99 (*35*)(Fig. 6, **lower panel**), would result in a futile cycle of continuous uptake of cholesterol and its mobilization from the PM to ER, resulting in the upregulation of MAM formation and activation, which in turn would cause the upregulation of SMases, ACAT1 activity, and LD deposition. Closing the cycle, this accumulation of cholesterol in the PM would also induce APP internalization and its interaction with, and cleavage by, BACE1 (*36*), and the downregulation of α-secretase activity (*37*). Interestingly, increases in exogenous cholesterol can mimic this scenario, resulting in APP internalization and C99 elevations (*38*), increases in the Aβ_42:40_ ratio (*39*), Tau phosphorylation (*40*), and hippocampal atrophy and cognitive impairment (*41*).

Our results are thus in support of a pathogenic role for C99 in AD. In the AD brain, there is a substantial increase in the production of amyloid, which originates from high levels of C99 fragments (*17*). Thus, the buildup of C99 in AD could be considered an early pathological hallmark that may elicit many of the molecular symptoms of the disease, including endosomal dysfunction (*16*), cognitive impairment, and hippocampal degeneration (*17*). Interestingly, others have found that changes in C99, rather than Aβ or AICD, could be behind some of the symptoms of dementia (*42*). In light of these reports and our own data, we believe that C99 toxicity in AD is mediated by its role on cholesterol metabolism.

Consistent with previous data (*29*), our results show that the buildup of C99 in MAM correlates with significant decreases in HMGCR and SREBP2 and the *de novo* synthesis of cholesterol, in an Aβ-and AICD-independent fashion. Thus, while the specific pathway needs to be elucidated, these data suggest that the accumulation of C99 prevents SREBP2 activation, impeding its function as a transcriptional factor (*6*). We note that one of the SREBP2-regulated genes is the LDL receptor, whose expression would also be reduced by SREBP2 downregulation in a feedback mechanism to control cholesterol levels (*6*). Paradoxically, our results also reveal that cholesterol uptake is highly induced in γ-secretase deficient cells. The intriguing failure of this negative feedback mechanism, suggests that this continuous cholesterol uptake might occur through one of the SREBP2-independent cholesterol receptors expressed in the cell (*43*).

As mentioned above, when in excess, a pool of “active” cholesterol in the PM will be proportionally transported to the ER cholesterol pool, where it will trigger feedback responses to maintain homeostasis (*7–9*). It has been suggested that this trafficking is regulated by cholesterol-sensing proteins and/or a specific cholesterol-sensing membrane domain in the ER associated with ACAT1 (*10*). Based on previous studies (*44*) and the results presented here, we propose that C99, via its CBD (*25*), acts as a cholesterol sensing protein, and that the MAM acts as signaling platform in the regulation of cholesterol homeostasis. Thus, it is possible that via this affinity domain, the accumulation of C99 generates cholesterol-rich areas needed for its cleavage by the γ-secretase complex. Hence, in the context of a deficient γ-secretase activity, uncut C99 will continue to recruit cholesterol to MAM, which helps explain the upregulation in MAM activity and ER-mitochondria connections found in cells from AD patients (*22, 24*). In agreement with this idea, C99^CBD^ defective in cholesterol binding failed to promote the upregulation of MAM functionality.

This effect on the formation and activation of lipid raft domains has been shown for the sigma-1 receptor, a MAM-resident protein (*45*) that, via its capacity to bind cholesterol, it causes the remodeling of lipid rafts and the regulation of the signaling molecules that are present in them (*46*).

Overall, we believe that our hypothesis suggests a potential mechanism to explain the seminal role of cholesterol in AD, underscored by the multiple genetic studies that have identified polymorphisms in genes related to cholesterol metabolism and the incidence of AD (*47*). Moreover, our data helps clarify the interdependence between cholesterol and APP metabolism (*38*); and the controversial association between cholesterol levels in AD (*48*), which we believe that could be rooted in the fact that defects in neuronal transmission could be caused by alterations in the distribution of subcellular cholesterol, rather than overall changes in cholesterol concentration.

Taken together, we propose a model in which C99 accumulation and increased cholesterol uptake occurs early in the pathogenesis of AD. Such a model would help create a framework to understand the role of cholesterol as both cause and consequence in the pathogenesis of AD, and the participation of many genetic loci associated with lipid metabolism, and specifically cholesterol regulation, in the pathogenesis of AD. In addition, it supports the idea that the APP C-terminal fragment acts as a cholesterol sensor protein in the membrane (*44*), whose cleavage regulate lipid homeostasis in the cell, coordinating the lipid composition of the PM and the intracellular lipid sensing platforms, namely MAM.

## ACKNOWLEDGEMENTS

We thank Dr. Xu (Sanford Burnham Institute) for the APP-DKO cells. We also thank Taekyung Yun and Maria Kaufman for technical assistance and the APP^V717I^ iPSCs. This work was supported by the U.S. National Institutes of Health (K01-AG045335 and R01-AG056387-01 to E.A.-G.), Alzheimer’s Association (AARF-19-614721 to J.M.), Alzheimer’s Disease Research Center (ADRC) pilot grant (to J.M.), the Michael J. Fox and the Leir Foundation (to C.G-L), the Henry and Marilyn Taub Foundation (A.A.S.), and the national defense science and engineering graduate fellowship (FA9550-11-C-0028 to R.R.A.)

## Author Contributions

E.A.-G. conceived the project, designed and interpreted most of the experiments. J.M. and E.A.-G. wrote the paper. M.P., D.L., J.M., C.G.-L, R.R.A. and K.R.V. performed most of the experiments and edited the manuscript. Y.X. performed the lipidomic analysis. SY.K. and A.S. generated APP^V717I^ homozygous knockin iPSC lines under the guidance of A.A.S., who also edited the manuscript.

## Author Information

The authors declare no competing financial interests. Correspondence and requests for materials should be addressed to E.A.-G.

---

**Supplementary Figure 1.**
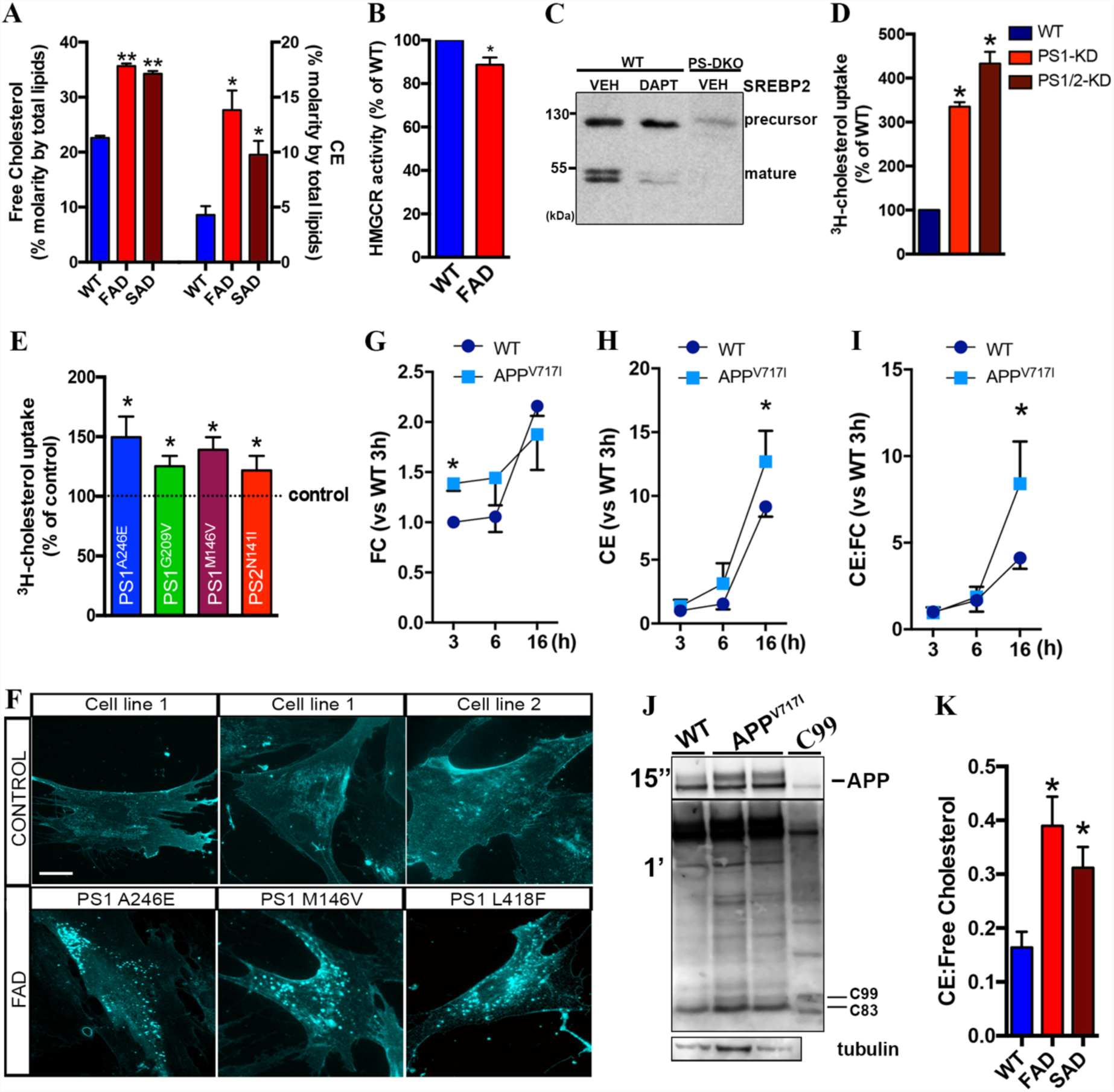
**(A)** Free cholesterol and cholesterol ester (CE) levels were measured in total homogenates of fibroblasts from FAD and SAD patients by lipidomics analysis (*, p<0.05, ** p<0.01, unpaired t-test vs WT; n = 4-8). **(B)** HMGCR activity was measured in cells from FAD patients and compared to control (WT) cells (* p<0.05, one-sample t-test vs WT, n = 3-4). **(C)** Measurement of SREBP2 levels by WB showed reduced levels of its mature/active form in DAPT-treated WT cells and a reduction of both the full-length/precursor and mature form in PS-DKO cells. **(D)** Incubation with ^3^H-cholesterol for 6h showed increased lipid uptake in neuroblastoma cell lines (Neuro-2a) where PS1 or PS1+PS2 had been transiently silenced (* p<0.05, one-sample t-test vs WT, n = 3). **(E)** Cholesterol uptake was measured by incorporation of ^3^H-cholesterol using different FAD cell lines. Dashed line indicates average WT/control levels. (* p<0.05, one-sample t-test vs WT; n=3). **(F)** Cholesterol distribution was detected by filipin staining of the indicated FAD fibroblasts and age-matched controls. Scale bar = 20 μm. **(G-H)** Cholesterol uptake and esterification was assessed by incubating APP^V717I^ and isogenic control cell lines with ^3^H-cholesterol and tracking its internalization and incorporation into ^3^H-cholesteryl esters (CE) by TLC upon different chase times (n=4). **(G)** Free cholesterol levels, FC. Two-way repeated measures ANOVA (Time 3&6h, Mutation): Mutation: F_(1,3)_ =14.36, p < 0.05, η=0.36. (* p<0.05 vs WT counterpart). **(H)** CE. Two-way repeated measures ANOVA (Time, Mutation): Time: F_(2,6)_=53.24, p < 0.001, η=0.76; Mutation: n.s.; Time × Mutation: F_(2,6)_=6.2, p < 0.05, η=0.017. (* p<0.05 vs WT counterpart). **(I)** The CE:FC ratio was calculated by lipidomics analysis. Two-way repeated measures ANOVA (Time, Mutation): Time: F_(2,6)_=14.38, p < 0.01, η=0.51; Mutation: n.s.; Time × Mutation: F_(2,6)_=5.2, p<0.05, η=0.09. (* p<0.05 vs WT counterpart). **(J)** C99 levels measured by WB in the APP^V717I^ and isogenic control cell lines. Total homogenates of C99-transfected APP-DKO cells were used as a control for C99 signal. APP is also shown. **(K)** Ratio of CE:free cholesterol measured by lipidomics of the indicated cells as in A. (*, p<0.05, unpaired t-test vs WT; n = 4-8).

**Supplementary Figure 2.**
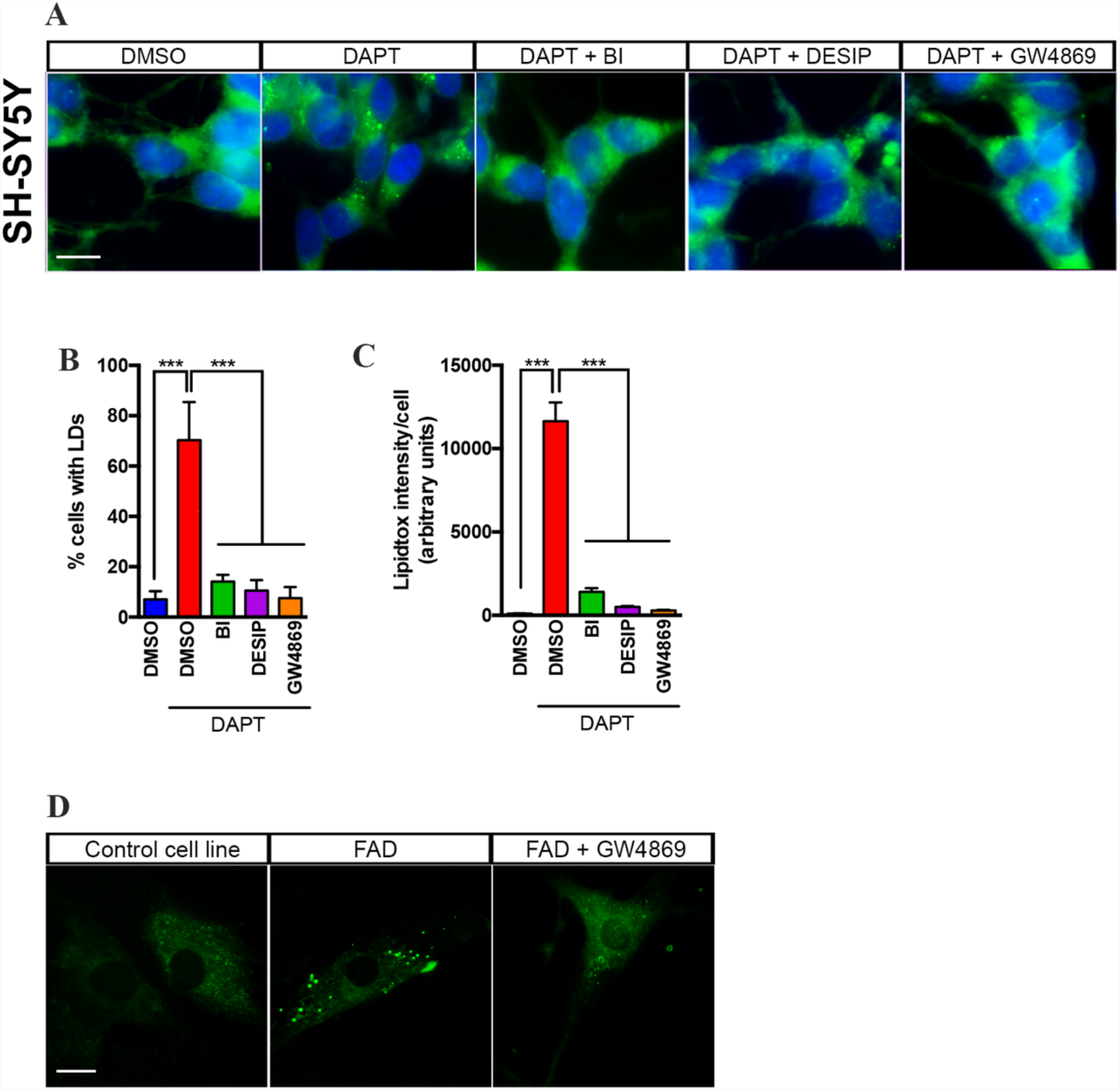
**(A-C)** Human neuroblastoma cells (SH-5YSY) were incubated with the indicated treatments for 12-16h or DMSO (VEH) and stained with LipidTox Green to detect lipid droplets upon different treatments. Scale bar = 20 μm. Note that treatment with the γ-secretase inhibitor, DAPT, caused an increase in either the % of cells with LDs **(B)** or the intensity of staining **(C)**, as quantified by ImageJ analysis. One-way ANOVA for B: F_(4,20)_=94.15, p < 0.001; for C: F_(4,20)_=65.11, p < 0.001 (n=5; *** p<0.001). **(D)** Fibroblasts from a FAD patient were incubated with SMase inhibitor (5 μM GW4869) for 12-16h and stained with Lipidtox Green to detect lipid droplets. Scale bar = 20 μm.

**Supplementary Figure 3.**
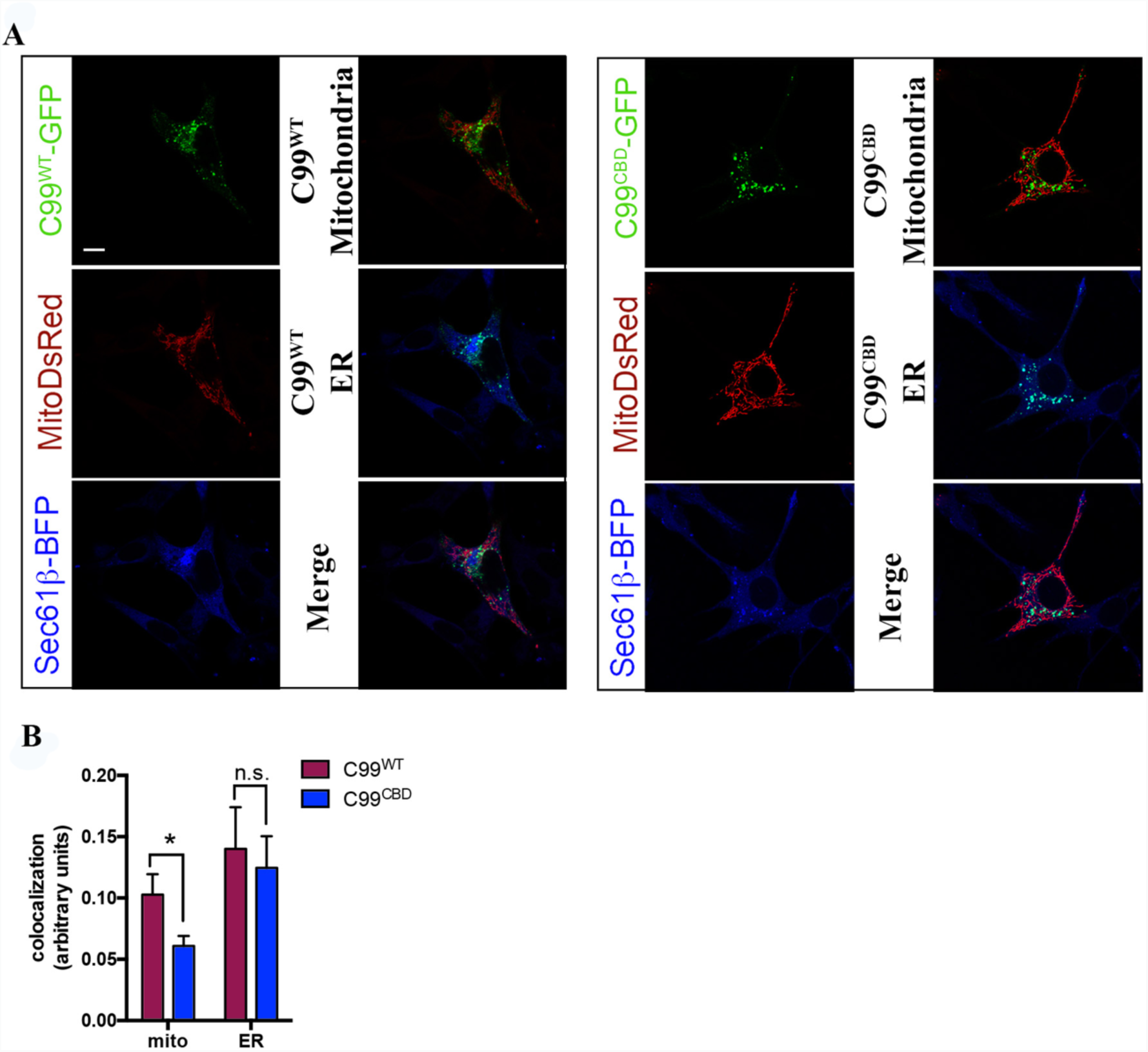
**(A)** Single green, red and blue confocal images show C99, MitoDsRed and Sec61β-BFP, respectively, as used in Fig. 3A. (**B**) Quantification of merged signals indicate decreased colocalization between C99^CBD^ and mitochondria, when compared to C99^WT^.

**Supplementary Figure 4.**
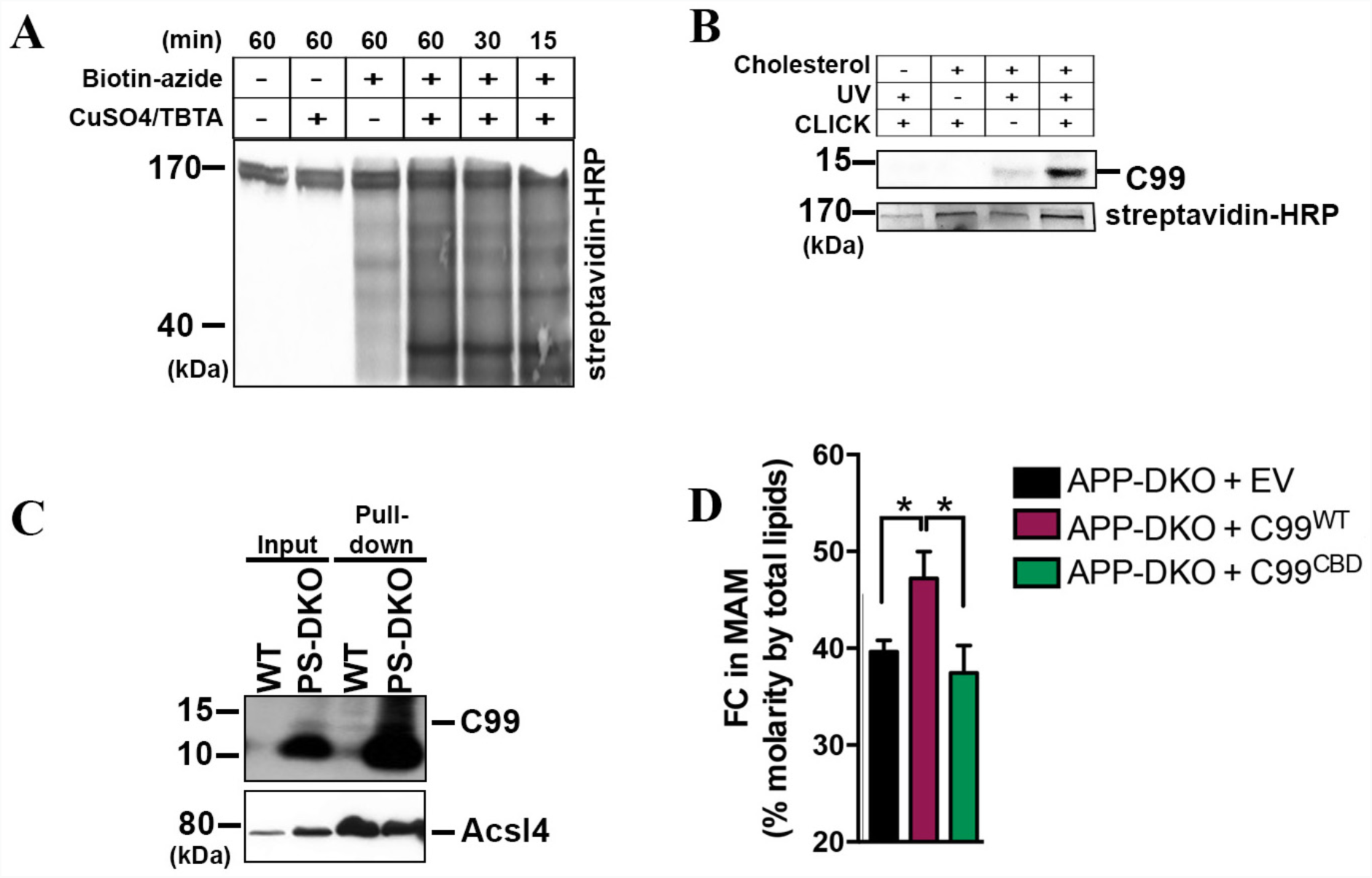
**(A)** Biotinylation levels were detected using streptavidin-HRP to optimize CLICK chemistry conditions. Note how an incubation of 15min is sufficient to obtain specific biotinylation. **(B)** C99 immunoblot of the pull-down following PhotoClick chemistry. Note that C99 levels in the pull-down are undetectable when either PhotoClick cholesterol, UV or CLICK chemistry is not used, supporting the specificity of the method. Streptavidin-HRP against endogenously biotinylated proteins was used as a loading control. **(C)** MAM fractions from WT or PS-DKO cells were subjected to PhotoClick chemistry and the input and pull-down were analyzed by western blot for C99 (A8717 antibody). **(D)** Levels of free cholesterol (FC) analyzed by lipidomics after subcellular fractionation to obtain MAM from the indicated cells. One-way ANOVA F_(2,9)_ = 4.59, p < 0.05. (n=4; * p<0.05).

**Supplementary Figure 5.**
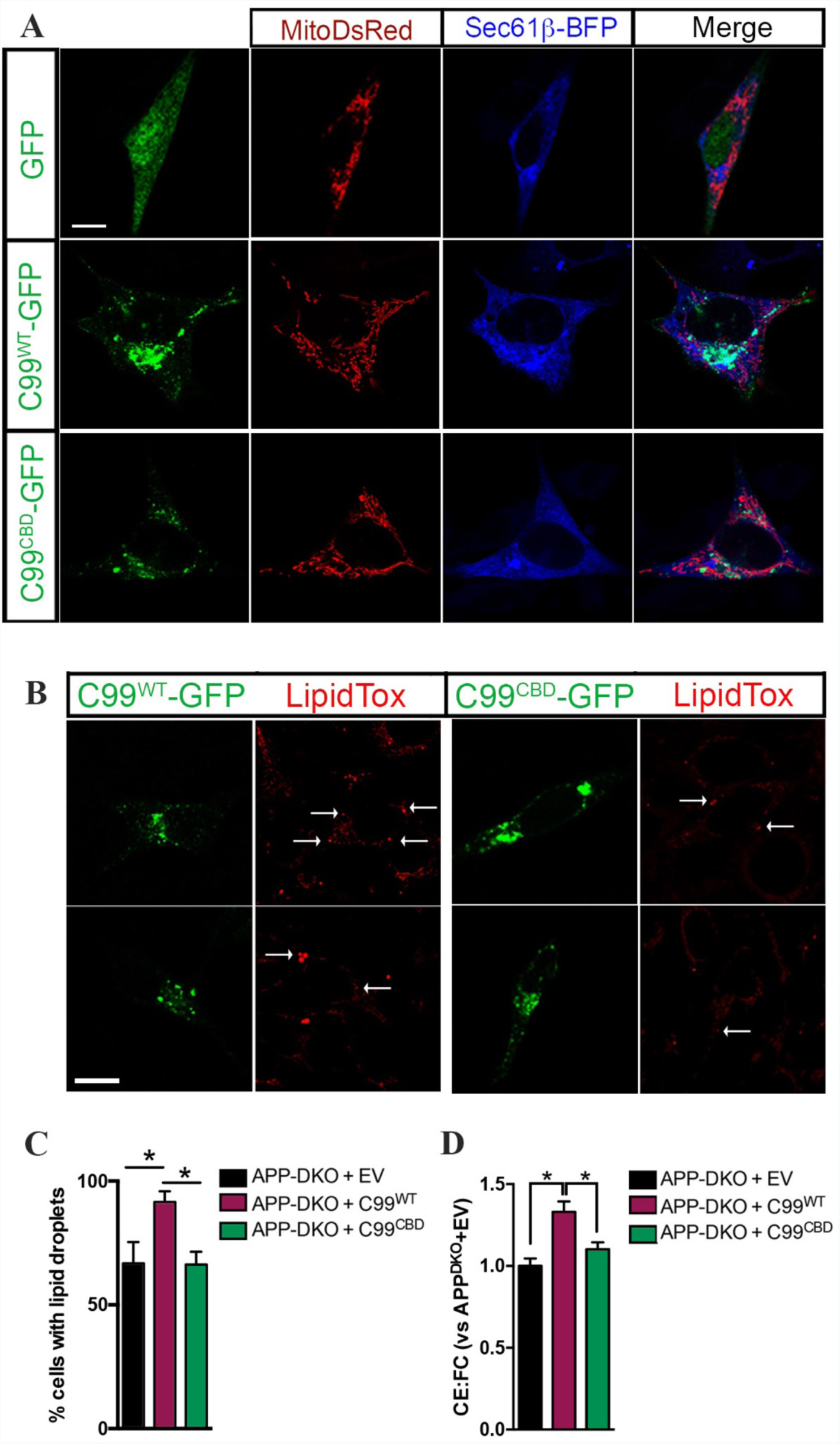
**(A)** Single green, red and blue confocal images show GFP/C99, MitoDsRed and Sec61β-BFP, respectively, as used in Fig. 5A. Merged images are also shown. **(B)** Representative images of APP-DKO cells expressing either GFP, C99^WT^-GFP or C99^CBD^-GFP and stained with LipidTox Red to detect LDs, marked with a white arrow. Scale bar = 15 μm. **(C)** The percentage of cells showing LDs was significantly different between groups. One-way ANOVA F_(2,6)_ = 5.25, p < 0.05 (30-50 cells/condition from at least 3 independent experiments; *p<0.05). EV, empty vector. **(D)** Ratio of cholesterol-esters: free cholesterol (CE:FC) of the indicated cells analyzed by lipidomics. One-way ANOVA F_(2,9)_ = 10.78, p < 0.01 (n=4; *p<0.05). EV, empty vector.

## REFERENCES

1. E. Lauwers, R. Goodchild, P. Verstreken, Membrane Lipids in Presynaptic Function and Disease. Neuron 90, 11–25 (2016).

2. D. Lingwood, K. Simons, Lipid rafts as a membrane-organizing principle. Science 327, 46–50 (2010).

3. R. F. Epand et al., Juxtamembrane protein segments that contribute to recruitment of cholesterol into domains. Biochemistry 45, 6105–6114 (2006).

4. T. Y. Chang, C. C. Chang, N. Ohgami, Y. Yamauchi, Cholesterol sensing, trafficking, and esterification. Annu Rev Cell Dev Biol 22, 129–157 (2006).

5. C. Yu, M. Alterman, R. T. Dobrowsky, Ceramide displaces cholesterol from lipid rafts and decreases the association of the cholesterol binding protein caveolin-1. J Lipid Res 46, 1678–1691 (2005).

6. M. S. Brown, J. L. Goldstein, A proteolytic pathway that controls the cholesterol content of membranes, cells, and blood. Proc Natl Acad Sci U S A 96, 11041–11048 (1999).

7. A. Das, M. S. Brown, D. D. Anderson, J. L. Goldstein, A. Radhakrishnan, Three pools of plasma membrane cholesterol and their relation to cholesterol homeostasis. Elife 3, (2014).

8. R. E. Infante, A. Radhakrishnan, Continuous transport of a small fraction of plasma membrane cholesterol to endoplasmic reticulum regulates total cellular cholesterol. Elife 6, (2017).

9. D. Y. Litvinov, E. V. Savushkin, A. D. Dergunov, Intracellular and Plasma Membrane Events in Cholesterol Transport and Homeostasis. J Lipids 2018, 3965054 (2018).

10. Y. Lange, J. Ye, M. Rigney, T. L. Steck, Regulation of endoplasmic reticulum cholesterol by plasma membrane cholesterol. J Lipid Res 40, 2264–2270 (1999).

11. A. M. Petrov, M. R. Kasimov, A. L. Zefirov, Brain Cholesterol Metabolism and Its Defects: Linkage to Neurodegenerative Diseases and Synaptic Dysfunction. Acta Naturae 8, 58–73 (2016).

12. G. Di Paolo, T. W. Kim, Linking lipids to AD: cholesterol and beyond. Nat Rev Neurosci 12, 284–296 (2011).

13. M. Goedert, M. G. Spillantini, A century of Alzheimer’s disease. Science 314, 777–781 (2006).

14. J. M. Cordy, N. M. Hooper, A. J. Turner, The involvement of lipid rafts in Alzheimer’s disease. Mol. Membr. Biol. 23, 111–122 (2006).

15. J. Shen, R. J. Kelleher, 3rd, The presenilin hypothesis of Alzheimer’s disease: evidence for a loss-of-function pathogenic mechanism. Proc. Natl. Acad. Sci. USA 104, 403–409 (2007).

16. Y. Jiang et al., Alzheimer’s-related endosome dysfunction in Down syndrome is Abeta-independent but requires APP and is reversed by BACE-1 inhibition. Proc Natl Acad Sci U S A 107, 1630–1635 (2010).

17. I. Lauritzen et al., The beta-secretase-derived C-terminal fragment of betaAPP, C99, but not Abeta, is a key contributor to early intraneuronal lesions in triple-transgenic mouse hippocampus. J Neurosci 32, 16243–16255a (2012).

18. B. Schreiner, L. Hedskog, B. Wiehager, M. Ankarcrona, Amyloid-beta peptides are generated in mitochondria-associated endoplasmic reticulum membranes. J Alzheimers Dis 43, 369–374 (2015).

19. E. Area-Gomez et al., Presenilins are enriched in endoplasmic reticulum membranes associated with mitochondria. Am. J. Pathol. 175, 1810–1816 (2009).

20. M. Pera et al., Increased localization of APP-C99 in mitochondria-associated ER membranes causes mitochondrial dysfunction in AD. EMBO J 36, 3356–3371 (2017).

21. T. Hayashi, M. Fujimoto, Detergent-resistant microdomains determine the localization of sigma-1 receptors to the endoplasmic reticulum-mitochondria junction. Mol. Pharmacol. 77, 517–528 (2010).

22. E. Area-Gomez et al., Upregulated function of mitochondria-associated ER membranes in AD. EMBO J, (2012).

23. J. E. Vance, MAM (mitochondria-associated membranes) in mammalian cells: Lipids and beyond. Biochim Biophys Acta 1841, 595–609 (2014).

24. L. Hedskog et al., Modulation of the endoplasmic reticulum-mitochondria interface in Alzheimer’s disease and related models. Proc Natl Acad Sci U S A 110, 7916–7921 (2013).

25. P. J. Barrett et al., The amyloid precursor protein has a flexible transmembrane domain and binds cholesterol. Science 336, 1168–1171 (2012).

26. A. Herreman et al., Total inactivation of γ-secretase activity in presenilin-deficient embryonic stem cells. Nat. Cell Biol. 2, 461–462 (2000).

27. R. B. Chan et al., Comparative lipidomic analysis of mouse and human brain with Alzheimer disease. J Biol Chem 287, 2678–2688 (2012).

28. X. Zhang et al., Hippocampal network oscillations in APP/APLP2-deficient mice. PLoS One 8, e61198 (2013).

29. N. Pierrot et al., Amyloid precursor protein controls cholesterol turnover needed for neuronal activity. EMBO Mol Med 5, 608–625 (2013).

30. J. P. Slotte, E. L. Bierman, Depletion of plasma-membrane sphingomyelin rapidly alters the distribution of cholesterol between plasma membranes and intracellular cholesterol pools in cultured fibroblasts. Biochem J 250, 653–658 (1988).

31. S. Endapally et al., Molecular Discrimination between Two Conformations of Sphingomyelin in Plasma Membranes. Cell 176, 1040–1053 e1017 (2019).

32. A. J. Beel, M. Sakakura, P. J. Barrett, C. R. Sanders, Direct binding of cholesterol to the amyloid precursor protein: An important interaction in lipid-Alzheimer’s disease relationships? Biochim. Biophys. Acta 1801, 975–982 (2010).

33. J. J. Hulce, A. B. Cognetta, S. E. Tully, B. F. Cravatt, Proteome-wide mapping of cholesterol-interacting proteins in mammalian cells. Nat Methods 10, 259–264 (2013).

34. M. Fujimoto, T. Hayashi, T. P. Su, The role of cholesterol in the association of endoplasmic reticulum membranes with mitochondria. Biochem Biophys Res Commun 417, 635–639 (2012).

35. J. Shen, R. J. Kelleher, 3rd, The presenilin hypothesis of Alzheimer’s disease: evidence for a loss-of-function pathogenic mechanism. Proc Natl Acad Sci U S A 104, 403–409 (2007).

36. J. C. Cossec et al., Clathrin-dependent APP endocytosis and Abeta secretion are highly sensitive to the level of plasma membrane cholesterol. Biochim Biophys Acta 1801, 846–852 (2010).

37. W. Wang et al., Amyloid precursor protein alpha- and beta-cleaved ectodomains exert opposing control of cholesterol homeostasis via SREBP2. FASEB J 28, 849–860 (2014).

38. C. Marquer et al., Local cholesterol increase triggers amyloid precursor protein-Bace1 clustering in lipid rafts and rapid endocytosis. FASEB J 25, 1295–1305 (2011).

39. C. Marquer et al., Increasing membrane cholesterol of neurons in culture recapitulates Alzheimer’s disease early phenotypes. Mol Neurodegener 9, 60 (2014).

40. R. van der Kant et al., Cholesterol Metabolism Is a Druggable Axis that Independently Regulates Tau and Amyloid-beta in iPSC-Derived AD Neurons. Cell Stem Cell 24, 363–375 e369 (2019).

41. F. Djelti et al., CYP46A1 inhibition, brain cholesterol accumulation and neurodegeneration pave the way for Alzheimer’s disease. Brain 138, 2383–2398 (2015).

42. R. Tamayev, S. Matsuda, L. D’Adamio, beta-but not gamma-secretase proteolysis of APP causes synaptic and memory deficits in a mouse model of dementia. EMBO Mol Med 4, 171–179 (2012).

43. R. S. Makar, P. E. Lipsky,, Multiple mechanisms, independent of sterol regulatory element binding proteins, regulate low density lipoprotein gene transcription. J Lipid Res 41, 762–774 (2000).

44. A. J. Beel et al., Structural studies of the transmembrane C-terminal domain of the amyloid precursor protein (APP): does APP function as a cholesterol sensor. Biochemistry 47, 9428–9446 (2008).

45. T. Hayashi, T. P. Su, Sigma-1 receptor chaperones at the ER-mitochondrion interface regulate Ca2^+^ signaling and cell survival. Cell 131, 596–610 (2007).

46. C. P. Palmer, R. Mahen, E. Schnell, M. B. Djamgoz, E. Aydar, Sigma-1 receptors bind cholesterol and remodel lipid rafts in breast cancer cell lines. Cancer Res 67, 11166–11175 (2007).

47. H. K. Dong, J. A. Gim, H. S. Kim, Integrated late onset Alzheimer’s disease (LOAD) susceptibility genes: Cholesterol metabolism and trafficking perspectives. Gene 597, 10–16 (2017).

48. W. G. Wood, L. Li, W. E. Muller, G. P. Eckert, Cholesterol as a causative factor in Alzheimer’s disease: a debatable hypothesis. J Neurochem 129, 559–572 (2014).

